# Reply to Barton et al: signatures of natural selection during the Black Death

**DOI:** 10.1101/2023.04.06.535944

**Authors:** Tauras P. Vilgalys, Jennifer Klunk, Christian E. Demeure, Xiaoheng Cheng, Mari Shiratori, Julien Madej, Rémi Beau, Derek Elli, Maria I. Patino, Rebecca Redfern, Sharon N. DeWitte, Julia A. Gamble, Jesper L. Boldsen, Ann Carmichael, Nükhet Varlik, Katherine Eaton, Jean-Christophe Grenier, G. Brian Golding, Alison Devault, Jean-Marie Rouillard, Vania Yotova, Renata Sindeaux, Chun Jimmie Ye, Matin Bikaran, Anne Dumaine, Jessica F Brinkworth, Dominique Missiakas, Guy A. Rouleau, Matthias Steinrücken, Javier Pizarro-Cerdá, Hendrik N. Poinar, Luis B. Barreiro

## Abstract

Barton *et al*.^1^ raise several statistical concerns regarding our original analyses^2^ that highlight the challenge of inferring natural selection using ancient genomic data. We show here that these concerns have limited impact on our original conclusions. Specifically, we recover the same signature of enrichment for high F_ST_ values at the immune loci relative to putatively neutral sites after switching the allele frequency estimation method to a maximum likelihood approach, filtering to only consider known human variants, and down-sampling our data to the same mean coverage across sites. Furthermore, using permutations, we show that the rs2549794 variant near *ERAP2* continues to emerge as the strongest candidate for selection (p = 1.2×10^−5^), falling below the Bonferroni-corrected significance threshold recommended by Barton *et al*. Importantly, the evidence for selection on *ERAP2* is further supported by functional data demonstrating the impact of the *ERAP2* genotype on the immune response to *Y. pestis* and by epidemiological data from an independent group showing that the putatively selected allele during the Black Death protects against severe respiratory infection in contemporary populations.

---

We thank Barton *et al*.^1^ for their careful consideration of our work^2^. They raise important concerns about our approach, leading them to conclude that our data set is underpowered to shed light on the evolution of the human genome in response to the Black Death. For example, they identify a source of bias in our method for allele frequency estimation. Below, we correct this bias in all analyses by using maximum-likelihood (ML)-based allele frequency estimates (as suggested by Barton *et al*.). They also raise legitimate concerns about how our relatively small sample size (n = 206) makes it challenging to definitively answer the question of how the Black Death may have shaped human evolution.

Ancient DNA studies are inherently limited by small sample sizes. Knowing that small samples are especially vulnerable to statistical bias, we carefully developed an integrative study design that combined multiple lines of evidence including sequence-based signatures of selection as well as functional experiments. Specifically, in our analysis, we combined a temporal sampling strategy of well dated ancestral remains with tight temporal constraints situated around the Black Death (before, during, and after the putative selective event), replication across two different population cohorts, contemporary evidence for balancing selection, and experimental data on the host cellular response to *Yersinia pestis* infection, to build a case for selection. Our functional data—which is extremely rare in ancient DNA studies—combined with epidemiological evidence from an independent group, strongly support the case for selection in *ERAP2*^3^ during the Black Death.

Despite the sum of our evidence, Barton *et al*.’s critique focuses almost entirely on the first stage of our study, which suggests selection on immune genes during the Black Death in the London cohort. They randomly permuted pre- and post-Black Death labels to show that there is a ∼7% probability of observing an enrichment of high F_ST_ values as large or larger than that observed in the London data — a result they attribute to differences in mean sequencing coverage between the immune loci and the putatively neutral loci in our data set. However, when considering both the London and Denmark cohorts together, we find permutations that yield higher F_ST_ values in *both* cohorts occur only 0.68% of the time (i.e., p = 0.0068). More generally, when using ML-based allele frequency estimates, which are not biased by sequencing coverage (Figure S1), and filtering to only consider known human variants, we found that the enrichment of high F_ST_ values at immune loci is similar to what we originally reported (Figure 1A). Additionally, after down-sampling our sequencing data to the same mean coverage across neutral and immune sites, we still recover a signature of enrichment for high F_ST_ values at the immune sites versus the putatively neutral sites, casting doubt on Barton *et al*.’s suggestion that coverage alone explains this enrichment (Figure 1B).

**Figure 1:**
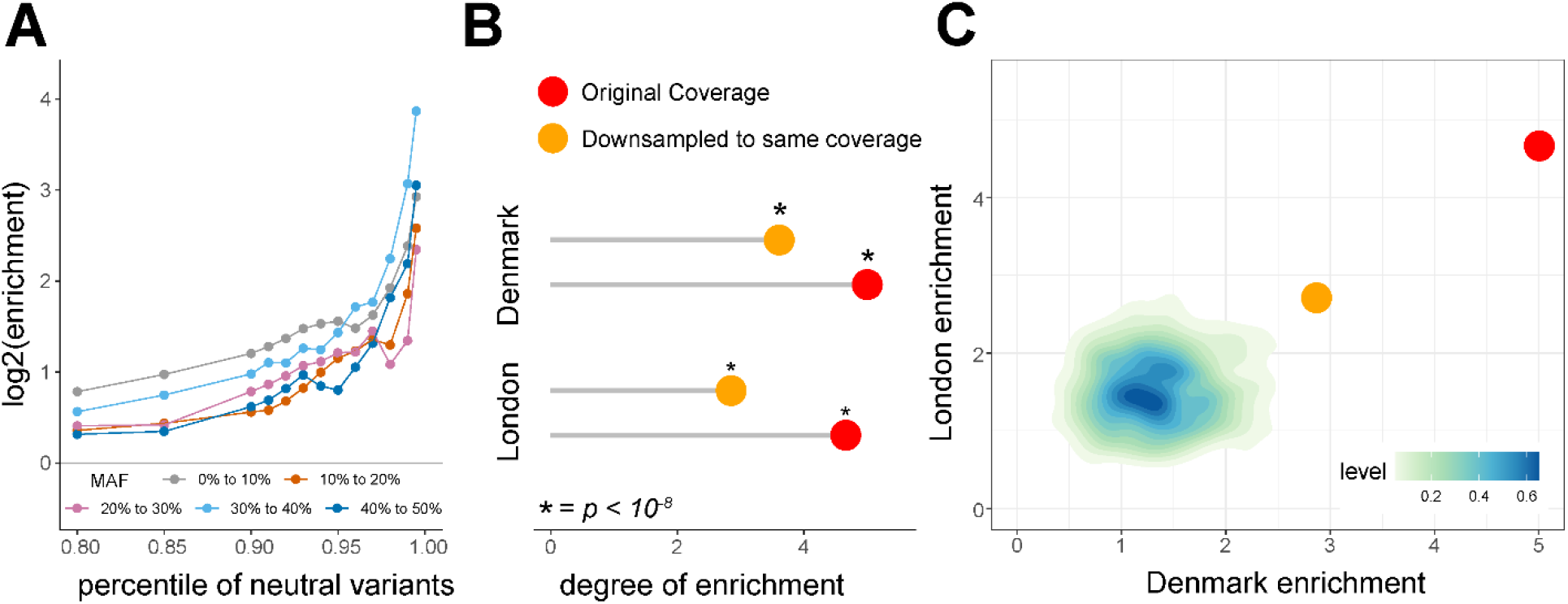
Enrichment of sites with high F_ST_ in candidate immune loci compared to putatively neutral loci is not an artifact of allele frequency estimation method or coverage. **(A)** Estimating allele frequencies using the ML method suggested by Barton *et al*. shows a significant enrichment of highly differentiated sites in immune loci relative to putatively neutral sites when comparing the pre-Black Death population to the post-Black Death population in London. **(B)** Down-sampling our data such that coverage was even across exon, immune, and neutral sites preserves the observed enrichments. The figure shows the degree of enrichment (odds-ratio) for candidate sites that exceed the 99^th^ percentile of neutral, MAF-matched F_ST_ values. Significance was assessed using a binomial test comparing the number of sites observed to pass the 99^th^ percentile to the number expected by chance. We note that down-sampling discards ∼1/3 of the data, which likely reduces our power to detect a significant enrichment of high F_ST_ in immune loci relative to putatively neutral sites. **(C)** Heat map of enrichment statistics obtained by permuting sites across locus type (neutral versus candidate immune loci). Observed enrichments for the down-sampled (orange) and full (red) datasets were larger than observed in any of the 10,000 permutations.

Additionally, we disagree that permutations across time (pre-versus post-Black Death samples) produce the appropriate null distribution for evaluating unusual patterns of evolution in immune genes. Our hypothesis was that the target immune genes would show greater divergence due to the Black Death than putatively neutral regions of the same genomes—a comparison that allows us to identify outlier genes and sites in the immune loci relative to other sites that experienced the same effects of population structure, migration, and other demographic processes. In contrast, the permutation-based null used by Barton *et al*. assumes that F_ST_ is 0 and all non-0 values simply reflect measurement noise. Even though the pre- and post-Black Death samples are close in time, this assumption does not appear to be reasonable: observed F_ST_ values from the neutral sites deviate from those generated through permutation (Figure S2). As an alternative approach, we permuted

SNPs in our sample between the “putatively neutral” and “immune” categories. Across 10,000 such permutations, none produced enrichment scores for immune loci (relative to neutral loci) as large as those in the observed data (i.e., p < 0.0001; Figure 1C). Importantly, this method again corroborates our original findings, independent of p-values from a binomial test or from differences in coverage depth (Figure 1C).

To identify individual sites that may have been the target of selection, our original approach focused on identifying F_ST_ outliers, a common practice in scans of natural selection in humans^4–6^. We believe this approach was helpful in narrowing the initial search space on SNPs of likely biological interest. Indeed, although Barton *et al*. rightly point out that we should have filtered our set of initial variants to those known in modern humans (which we have done for all analyses reported here, in addition to our original measures to account for DNA damage), it is worth noting that 238 out of the 245 F_ST_ outlier SNPs discovered in the London cohort based on F_ST_ are known SNPs reported by the 1000 Genomes’ project^6^. Requiring independence evidence from two populations, in addition to making directional predictions about how allele frequencies should change across the Black Death pandemic, further increased the burden of evidence required to identify candidate targets for selection.

Barton *et al*. also suggest that our analysis should explicitly focus on p-value estimates for individual SNPs, followed by Bonferroni correction. While this was not our original approach, here we follow their suggestion and calculate site-specific p-values obtained after comparing observed F_ST_ values with values derived from permuting individuals between time points (pre-, during, and post-Black Death). Following our original criteria, we limited our analyses to sites where (i) in London, the direction of change from the pre-Black Death samples to those collected during the Black Death was opposite to the direction of change from pre-Black Death to post-Black Death samples; and (ii) the pre- to post-Black Death change in allele frequencies is in the same direction in London and Denmark (samples were not available during the Black Death for Denmark). rs2549794 near *ERAP2* remains the top hit among all variants (Figure 2, p = 1.2×10^−5^). Importantly, the p value for rs2549794 is lower than the conservative Bonferroni-corrected significance threshold proposed by Barton *et al*.: 0.05/1,758 = 2.8×10^−5^. Here, 1,758 corresponds to the total number of immune SNPs in our dataset with a MAF >5% and not only those that meet the criteria described above (limiting to known SNPs as suggested by Barton *et al*.). Another variant (rs2548527) in strong LD with rs2549794 (r^2^ = 0.79 in 1000Genomes’ GBR population) exhibits qualitatively similar changes in allele frequency between pre- and post-Black Death samples, indicating that our results for *ERAP2* are not due to potential DNA damage (Figure 2B). Notably, in a recent study on ancient DNA samples from Cambridgeshire, England, the putatively protective allele of *ERAP2* also increased in frequency from 52% pre-Black Death to 60% post-Black Death^7^. While Barton *et al*. highlight that Hui *et al*.^7^ do not replicate this SNP as an F_ST_ outlier, we note that this analysis only focused on only n=50 samples (approximately a third of our sample size for London only).

**Figure 2:**
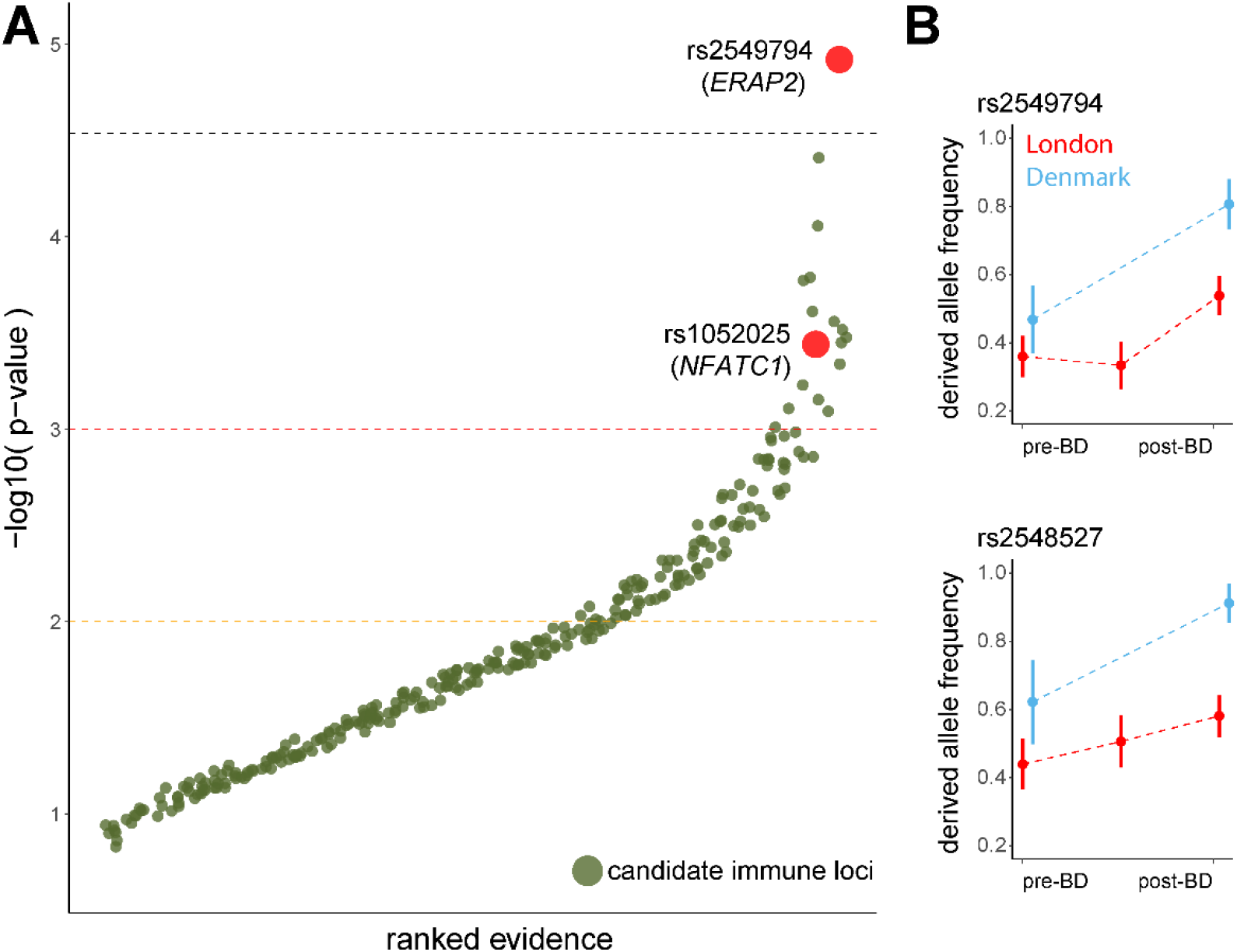
Loci ranked by evidence for positive selection, shown on the y-axis as the -log_10_ proportion of permutations where the F_ST_ was greater than the observed F_ST_ in both London and Denmark, the allele frequency change between pre- and post-Black Death was in the same direction for both London and Denmark, and the allele frequency change between pre- and during-Black Death in London was in the opposite direction. Candidate immune loci (n=290) are shown in green, and values are jittered along the x-axis to limit overlapping points. Dashed lines correspond to p-values of 0.01 (orange), 0.001 (red), and a Bonferroni corrected p-value threshold of 2.8×10^−5^ (black). Among our 4 original candidate loci, rs2549794 (*ERAP2*, p = 1.2×10^−5^) and rs1052025 (*NFATC1*, p = 3.6×10^−4^) are shown in red. The other two variants failed to meet the criteria where the changes in allele frequency between pre- vs post-Black Death and pre- vs during-Black Death should be in the opposite direction. That said, the levels of genetic differentiation observed in London remains unlikely by chance alone (rs11571319/*CTLA4*: p = 0.0608; rs17473484/*TICAM*: p = 0.0997). **(B)** Allele frequencies over time for rs2549794 and rs2548527, which are strongly linked and near *ERAP2*. Error bars represent the standard deviation based on bootstrapping individuals from that population and each time point 10,000 times. Allele frequencies for London are shown in red and for Denmark are shown in blue.

Finally, Barton *et al*. report that previous scans of selection^8–10^ do not support selection on *ERAP2*. This does not contradict our findings, since the methods used in those scans were designed to detect directional selection over many generations, which is not the scenario that we proposed for *ERAP2*. As discussed in our original manuscript, and supported by other studies^3,11,12^, we propose that the *ERAP2* variant has evolved under long-term balancing selection, with a short pulse of strong directional selection in response to the Black Death. Specifically, we suggest that the rs2549794-C allele was advantageous during the Black Death, but that it is also associated with a fitness cost that maintains a long-term balanced allele frequency. Importantly, the protective nature of this allele against the Black Death is supported by our functional data: we show that macrophages from individuals with the C allele engage a unique cytokine response and are better able to control bacterial growth upon infection with *Y. pestis*, compared to individuals with the T allele. Furthermore, we note that the selection coefficient we estimated for rs2549794 near ERAP2 cannot be directly compared to coefficients estimated for loci such as the lactase persistence allele or other cases of long-term selection. In our study, we modeled directional selective pressure during a short, 3-generation pulse, whereas, for example, selection for lactase persistence occurred over at least 150 generations^8^. In addition, as originally noted^2^, the estimate is associated with large uncertainty and therefore should be interpreted cautiously. More generally, we acknowledge the possibility that the selection coefficient we presented for rs2549794 may be over-estimated due to small sample sizes and ascertainment bias towards high F_ST_ variants, and quantifying this will require further investigation.

## Concluding remarks

In summary, the new analyses we present here reinforce our original conclusions; namely, that changes in allele frequencies at immune genes during the Black Death are larger than expected based on putatively neutral regions; that the temporal patterns of genetic differentiation at the *ERAP2* locus appear particularly unlikely under neutrality; and that *ERAP2* genotype has strong effects on *ERAP2* gene expression and the ability to restrict *Y. pestis* in human macrophages. These conclusions are largely unaffected by the issues highlighted by Barton *et al*.

Indeed, for classic cases of positive selection such as alleles for lactase persistence or malaria resistance (e.g., *G6PD* or the Duffy null allele), we argue that it is the *combination* of sequence-based evidence with functional, organismal, and epidemiological data that makes these cases of selection compelling. These are all crucial sources of evidence to support natural selection. Notably, recent evidence from an independent group shows that the variant we predict to have been protective during the Black Death is indeed protective against infectious diseases of bacterial origin, while also increasing the risk for Crohn’s disease and diabetes^3^. Such convergent evidence is indispensable given the constraints of ancient DNA studies, where achieving large sample sizes remains a challenge. Meanwhile, we look forward to future ancient DNA studies with larger sample sizes, and expanding to the resolution of entire genomes, to fully delineate the impact of the Black Death on the evolution of the human immune system.

## Methods

### Site curation and allele frequency estimates

Barton *et al*. raise a valid concern that our original dataset included many sites not previously identified as polymorphic in humans, many of which may have been introduced by DNA damage. We correct for that oversight by limiting our analyses here to variants that are also reported in the 1000G project^6^. Doing so results in a transition to transversion ratio of 3.18 and a strong correlation in allele frequency between our population in London and the 1000 Genomes project GBR population^6^ (r^2^ = 0.94, p < 10^−300^). As with all ancient DNA studies, we cannot completely rule out damage as a contributing factor to our genotype calls. However, as described in the original publication, we took several steps to minimize the impact of deamination and other forms of ancient DNA damage in our estimates of allele frequencies. Quality scores in each bam file were adjusted for DNA degradation based on their initial qualities, position in reads and damage patterns. In addition, we systematically trimmed off the first and last 4 bases of each sequencing read where the bulk of damage accrued for our samples is concentrated (Figure S1 of original publication). Moreover, we note that the average coverage across our most highly differentiated sites is considerably high for ancient DNA data (mean of 4.5x across all variants initially reported in Table S4, and 9.4x rs2549794 near *ERAP2)*, which minimizes the potential contribution of DNA damage to our genotype calls.

We also revise our allele frequency estimator to use the ML-based approach suggested by Barton *et al*. Specifically, we let the genotype likelihoods for individual *i* in 1:n be given by 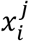 where *j* is the genotype denoted as 0, 1, or 2 alternate alleles. The ML estimate of the allele frequency *p* is then:

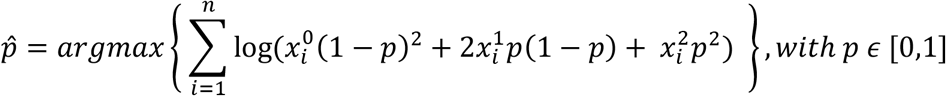

Prior to estimating F_ST_, We applied the numerical implementation of this estimator used by Barton *et al*. to estimate allele frequencies in each population and each time point, based on the genotype likelihoods reported in our original manuscript.

### Test for enrichment at immune loci

We replicated our previous analysis demonstrating an enrichment for highly differentiated candidate sites using these ML-based allele frequency estimates. Specifically, we identified highly differentiated sites by comparing the observed F_ST_ values for candidate immune sites against 200 MAF-matched neutral sites. Matched neutral sites were selected as the sites with closest MAF to the candidate variant. This represents a slight departure from our previous approach using MAF bins containing at least 200 neutral sites, which was necessary due to the fact that we have now limited our analyses to a smaller set of variants known to be polymorphic in humans.

### Coverage alone does not drive the observed enrichment

Barton *et al*. argue that differences in coverage between sets of sites drives the observed enrichment for high F_ST_ values among immune loci. To precisely test this hypothesis, we down-sampled our dataset such that the coverage was the same between the three capture sets (exons, GWAS, and neutral loci). Specifically, we estimated the coverage of each sample at analyzed sites using “samtools depth” and then used the “samtools view” with the -s option to down-sample each capture to the same, minimum coverage for that individual^13^. For samples where one of the capture sets was not used, we instead down-sampled sequence data sets for that sample to the average coverage for all individuals. We then estimated allele frequencies and calculated the enrichment of highly differentiated loci, as described above. As expected, this procedure resulted in similar coverage between the sets of immune and neutral loci (mean coverage at GWAS loci 5.416 ± 3.56x, at exonic loci 5.419 ± 3.57x, and at neutral loci 5.417 ± 3.58x).

### Permutation test for enrichment at immune loci

We first used the permutation approach employed by Barton *et al*. to test whether the observed enrichment of large F_ST_ values at candidate loci exceeded random expectations. Specifically, within each population, we randomly sampled individuals with replacement for each time point. Where Barton *et al*. only performed this analysis for the London pre-Black Death and London post-Black Death samples, our approach also considered the London samples collected during the Black Death and the pre-Black Death and post-Black Death samples collected in Denmark. We chose to perform sampling without replacement because, for the London cohort, we noticed that dependence across time-points resulted in strongly correlated estimates for the difference between pre- and post-BD and between pre- and during-BD time points. Running permutations without replacement does not impact our conclusions.

In addition, we used a second permutation approach where we permuted SNP assignment into the “putatively neutral” versus “immune” categories, while keeping the sizes of both categories identical to those in the original data. We repeated this process 10,000 times for each permutation approach, and for each iteration calculated the proportion of sites that exceeded 99% of neutral, MAF-matched loci. We then estimated the proportion of permutations in which more sites passed this threshold in the permuted data than in the observed data (which was none of the 10,000 permutations).

### Permutation test for enrichment at immune loci

Inspired by the permutation approach proposed by Barton *et al*., we next used permutations to calculate a site-specific, empirical p-value for each site that is independent of MAF-matched neutral loci. Following the same reasoning as in the original publication, we filtered for sites where (i) in London, the direction of change from the pre-Black Death samples to those collected during the Black Death was opposite to the direction of change between the pre-Black Death to post-Black Death samples; and (ii) the pre to post-Black Death allele frequencies change in the same direction in London and Denmark. We then calculated an empirical p-value as the proportion of one million permutations for which these conditions were also met and the permuted F_ST_ value exceeded the observed F_ST_ value in both London and Denmark.

### Allele frequencies and differentiation at rs2549794 and rs2548527

To show the change in allele frequency over time at variants near *ERAP2* (rs2549794 and rs2548527), we estimated the variance of the allele frequencies at each time point using ML allele frequency estimates by bootstrapping with 10,000 replicates. We plot the mean and standard deviation. Mean coverage of rs2549794 was more than twice that of rs2548527 (9.4x vs 3.7x) and many more individuals were missing genotype calls for rs2548527 (56 samples vs 6 for rs2549794), reducing confidence in the estimated allele frequencies at rs2548527.

## Data availability

No new data were generated for this work.

## Code availability

Code to replicate the analyses described here is available at

https://github.com/TaurVil/VilgalysKlunk_response_to_commentary

## Acknowledgements

We thank all members of the Barreiro lab and the Poinar lab for their constructive comments and feedback. We thank Jeremy Berg, Ran Blekhman, Yoav Gilad, Natalia Gonzales, Etienne Patin, George Perry, Lluis Quintana-Murci, and Jenny Tung for their comments on the manuscript. Computational resources were provided by the University of Chicago Research Computing Center. This work was supported by grant R01-GM134376 to LBB, HP, and JP-C, a grant from the Wenner-Gren Foundation to JB (8702), and the UChicago DDRCC, Center for Interdisciplinary Study of Inflammatory Intestinal Disorders (C-IID) (NIDDK P30 DK042086). HNP was supported by an Insight Grant #20008499 from the Social Sciences and Humanities Research Council of Canada (SSHRC) and The Canadian Institute for Advanced Research under the Humans and the Microbiome program. TPV was supported by NIH F32GM140568. XC and MSt were supported by grant R01GM146051.

## Competing interest declaration

JK, ADe, and J-MR declare financial interest in Daicel Arbor Biosciences, who provided the myBaits hybridization capture kits for this work. All other authors declare no competing interests.

**Figure S1:**
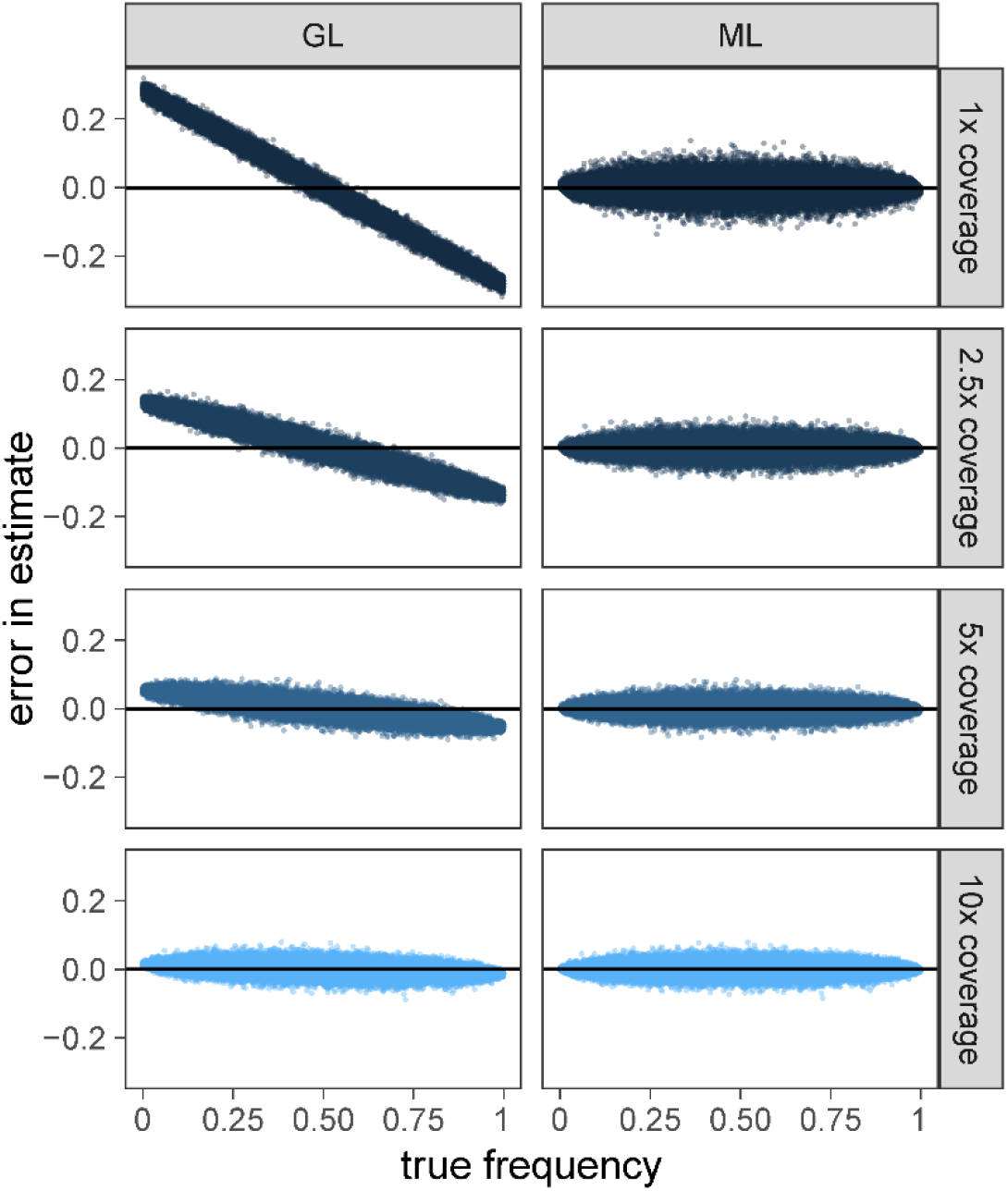
The difference between simulated “true” allele frequencies and the allele frequency estimated using either the genotype likelihoods (GL) in Klunk *et al*. or the maximum likelihood (ML) approach outlined by Barton *et al*. The GL approach shows a bias towards overestimating the frequency of rare variants, which is exaggerated at lower coverages. No similar bias is apparent using the ML approach.

**Figure S2:**
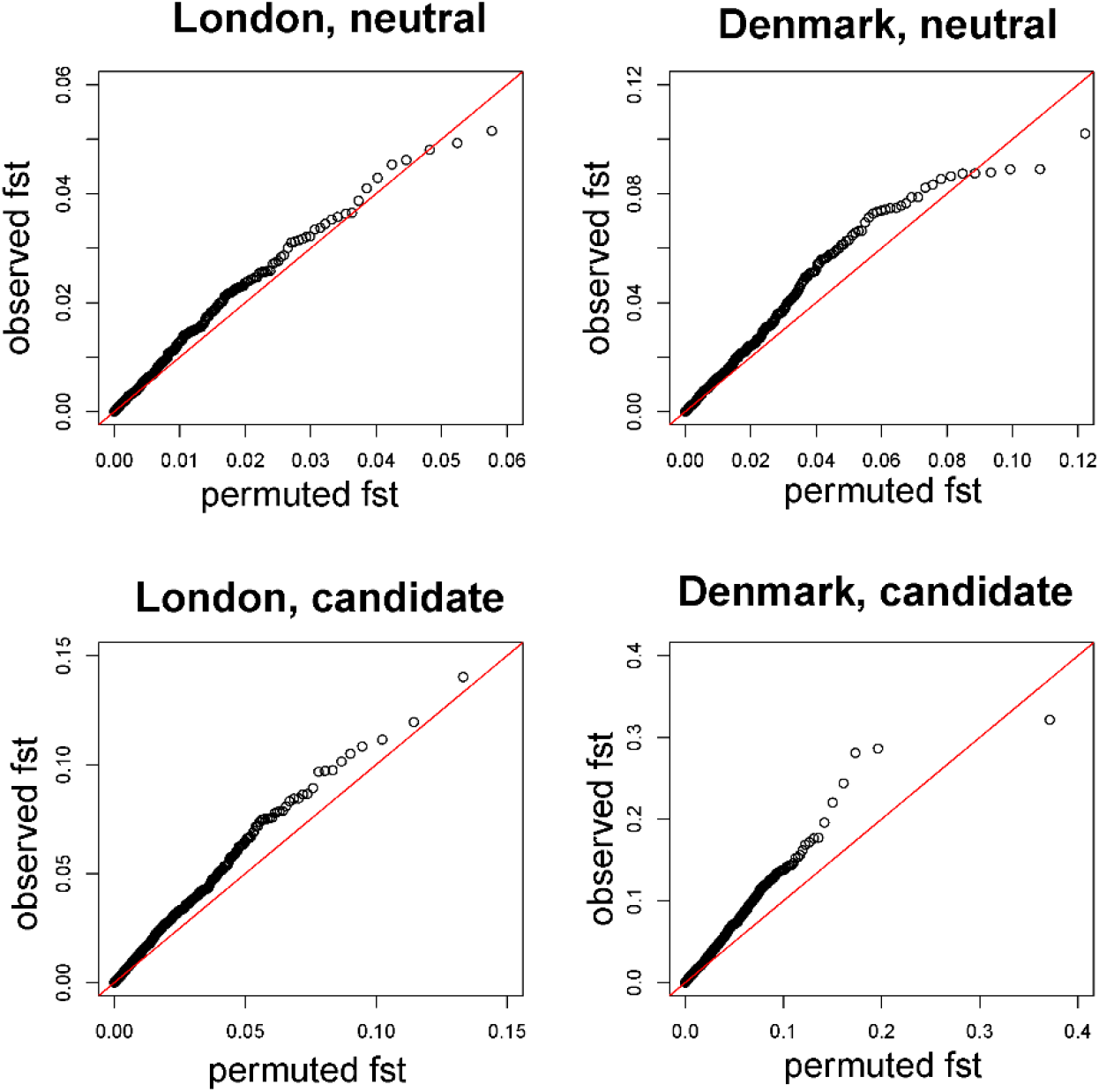
QQ-plots comparing the observed distribution of F_ST_ values to the distribution of F_ST_ values where individuals were randomized across time points. Observed data for both neutral and candidate loci show an enrichment of elevated F_ST_ values. Specifically, observed neutral F_ST_ values are elevated compared to those from permuted data (London p = 0.052; Denmark p = 0.0058).

